# CRTC, not phosphorylated CREB1, drives cAMP-induced transcription across diverse cell types

**DOI:** 10.64898/2026.07.27.739364

**Authors:** Charles H. Adelmann, Michelle Liu, Allison E. Greuel, Lingjuan Huang, Judith R. Boozer, Avanthika Venkatachalam, Xunwei Wu, Eleanor Ziarnik, Jiayin Tang, Ana Alcantara, Joshua P. Carreras, Suyeon Ryu, Charles Shi, Long Huynh, Tristan Vornbäumen, Garyfallia Papaioannou, David S.B. Hoon, Nir Hacohen, Marc N. Wein, David E. Fisher

**Affiliations:** Cutaneous Biology Research Center, Department of Dermatology, Massachusetts General Hospital, Boston, MA 02114, USA; Krantz Family Center for Cancer Research, Massachusetts General Hospital Cancer Center, Massachusetts General Hospital, Boston, MA 02114, USA; Department of Translational Molecular Medicine and the Genomic Sequencing Center, Saint John’s Cancer Institute, Santa Monica, CA 90404, USA; Department of Cancer Immunology and Virology, Dana-Farber Cancer Institute, Boston, MA 02115, USA; Department of Immunology, Harvard Medical School, Boston, MA 02115, USA; Department of Neurobiology, Harvard Medical School, Boston, MA 02115, USA; Endocrine Unit, Massachusetts General Hospital, Boston, MA 02114, USA; NGIH, Los Angeles, CA 90025, USA; Harvard Medical School, Boston, MA 02115, USA; Broad Institute of MIT and Harvard, Cambridge, MA 02142, USA

## Abstract

Specialized cell types layer cell-type-restricted proteins, metabolites, and organelles onto a shared foundation of core cellular processes. How this ubiquitous machinery generates lineage-specific outputs is central to both cell biology and therapeutic development. G-protein-coupled receptors (GPCRs) respond to receptor-restricted ligands to drive cell-type-specific transcription through cAMP-mediated activation of protein kinase A (PKA), which activates the transcription factor CREB1 through two parallel modes: direct phosphorylation at serine 133, and inhibition of salt-inducible kinases (SIK) that restrain the CRTC coactivators. How these modes integrate and contribute to signaling across cell types has remained unresolved, obscured by genetic redundancy and essentiality. Here we combine focused genetic analyses with a cross-lineage transcriptomic survey to dissect these parallel inputs. In *Creb1*/*Atf1*/*Crem* triple-knockout cells, a non-phosphorylatable mutant CREB1^S133A^ fully rescued endogenous target gene activation, while *Crtc1/Crtc2/Crtc3* ablation abolished transcription even with intact CREB1 serine 133 phosphorylation. As part of this mechanism, we found the annotated repressor ICER can instead act as a positive regulator, substituting for full-length CREB1 paralogs to drive a feedforward loop. Across melanocytes, hepatocytes, osteocytes, macrophages, and neurons, SIK inhibition recapitulated cAMP-PKA-driven transcription across both shared and cell-type-specific gene expression programs, with neurons a notable exception. These results invert the canonical model, placing CRTC recruitment as the dominant driver of CREB1-mediated transcription across diverse lineages, reframing how cAMP-PKA signaling can be interpreted and therapeutically targeted.

## Introduction

Cells of different types respond to distinct signals with distinct transcriptional programs, yet how shared signaling machinery produces these cell-type-specific outputs remains a fundamental question in cell biology. The PKA-CREB1 pathway exemplifies this challenge: the same pathway operates downstream of a broad range of G-protein-coupled receptors (GPCRs) across many different cell types to drive distinct transcriptional programs ^1,2^. For example, in melanocytes, the GPCR MC1R responds to the UV-induced ligand α-MSH to activate PKA-CREB1 and drive the transcription of *MITF*, a central regulator of pigmentation genes (Figure 1A)^3–6^, whereas in hepatocytes, the GCGR binds glucagon, secreted in response to low blood glucose, to activate PKA-CREB1 and drive the transcription of gluconeogenic enzymes such as *PCK1* and *G6PC1*^7,8^.

**Figure 1.**
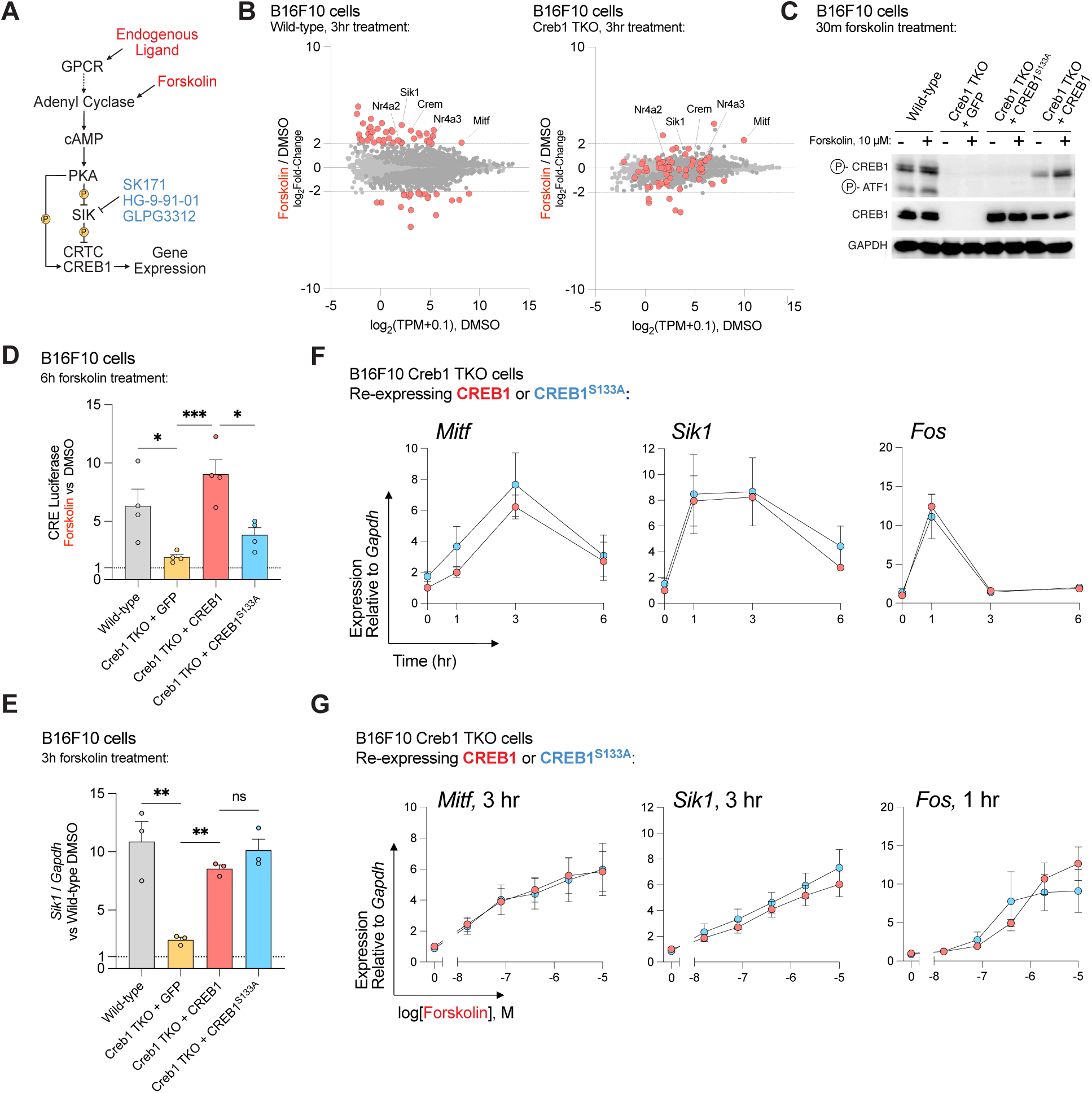
CREB1-controlled gene expression does not require CREB1 serine 133 phosphorylation. **(A)** Model of GPCR-induced transcription in melanocytes highlighting pharmacological tools used to activate the CRTC-CREB1 axis. **(B)** MA plot of transcriptional profiles of wild-type and Creb1 TKO cells following 3hr of 10 µM forskolin treatment . Dark gray points indicate *padj* < 0.01. Red points highlight genes that were strongly regulated in wild-type cells (|LFC| > 2, padj < 0.01). Analysis was restricted to protein-coding mRNAs, fold-change and adjusted p-values were calculated using DESeq2. **(C)** Immunoblot of wild-type, Creb1 TKO cells, and Creb1 TKO cells re-expressing wild-type human CREB1 or CREB1^S133A^ stimulated with DMSO or forskolin (10 µM). **(D)** CRE-driven luciferase activity from a transfected minimal reporter (4xCRE:HSV promoter) following 6 hr of forskolin (10 µM) treatment. Values were normalized to co-transfected constitutive Renilla and to DMSO-treated samples of matched genotype (n = 4). **(E)** Model CRTC-CREB1 target gene *Sik1* measured by RT-qPCR in Creb1 TKO cells re-expressing CREB1 or CREB1^S133A^, stimulated with DMSO or forskolin (10 µM) for 3 hr. Samples were normalized to *Gapdh* and to wild-type DMSO-treated cells. Baseline expression was unchanged across genotypes (Supplemental Figure 1B; n = 3). **(F)** Time-course analysis of *Mitf, Sik1,* and *Fos* induction measured by RT-qPCR in cells re-expressing CREB1 or CREB1^S133A^. Samples were normalized to *Gapdh* and to untreated wild-type CREB1 samples (n ≥ 4). **(G)** Dose-response analysis of *Mitf, Sik1*, and *Fos* induction across a ∼3-log forskolin concentration range (16 nM – 10 µM). *Mitf* and *Sik1* assays were measured after 3 hr; *Fos* after 1 hr. Samples were normalized to *Gapdh* and untreated CREB1 wild-type samples (n ≥ 5). *p<0.05, **p<0.01, ***p<0.001. Bar plot statistical tests between conditions were assessed by one-way ANOVA with Sidak’s multiple comparison test, while comparisons to theoretical mean (i.e., 1.0) was performed by one-sample Students t-test. Error bars are standard error of the mean.

GPCR activation stimulates adenylyl cyclase to produce cAMP, which activates PKA. Activated PKA in turn activates transcription through two parallel outputs that converge on the transcription factor CREB1 and its close paralogs ATF1 and CREM^1,2^. The first output is direct PKA-mediated phosphorylation of CREB1 on serine 133 within the kinase-inducible domain (KID) of its transactivation domain, an event that facilitates the recruitment of the transcriptional coactivators p300/CBP^1,9–11^. PKA-mediated phosphorylation of CREB1 on serine 133 is conserved across metazoans, including in hydra, nematodes and flies^12^. It has been extensively characterized and is the textbook^13^ explanation for PKA-driven CREB1-controlled transcription. The second output runs through the salt-inducible kinases (SIK1, SIK2, and SIK3). Here, PKA phosphorylation inhibits SIK activity, which otherwise results in the phosphorylation of the CRTC transcriptional coactivators (CRTC1, CRTC2, and CRTC3) to retain them in the cytosol through phosphorylation-dependent 14-3-3 protein binding^2,14–17^. Upon PKA-mediated SIK inhibition, CRTCs become dephosphorylated, translocate to the nucleus, bind CREB1, and potentiate transcription through their own transactivation domain^18^.

The relative contribution of CREB1 phosphorylation versus CRTC activation via SIK inhibition in specific biological systems is unknown. Systematic dissection of the PKA-CREB1 pathway has been complicated by the two- to three-member redundancy of every component and by the essentiality of these components in both animals and cell lines. This question has gained particular urgency with the recent development of SIK inhibitors as therapeutics, which have shown promise in preclinical models of ulcerative colitis, psoriasis, rheumatoid arthritis, osteoporosis, fracture healing, skin pigmentation, and various cancers^5,19–22^, while genetic analysis also implicates SIK as a key mediator of obesity and sleep regulation^23–26^. Unlike upstream GPCR agonists, which engage both arms of the pathway, SIK inhibitors act downstream of PKA and bypass CREB1 serine 133 phosphorylation entirely. Predicting their effects across target and non-target tissues therefore requires knowing how much of PKA-driven transcription depends on the SIK–CRTC arm in each cell type.

Here, we sought to define the relative contributions of the two PKA-controlled inputs into CREB1, focusing on both the fundamental mechanisms of the pathway and the functional consequences of small molecule SIK inhibition. Through genetic analysis, we established that CRTCs are necessary for transcription of most PKA-regulated targets, while PKA-mediated CREB1 serine 133 phosphorylation is largely dispensable. This insight led us to discover that ICER1, a previously annotated repressor transcribed from the *CREM* locus, can act as a positive regulator of PKA-CREB1 signaling, and it functions as a feedforward amplifier of pigmentation genes in the melanocyte lineage. Consistent with the dominance of the SIK-CRTC arm, a transcriptomic survey of primary and tumor cell lines from diverse lineages found that chemical SIK inhibition closely recapitulates most of the cAMP-PKA-regulated transcriptional response across cell types. Together, our findings systematically define the contributions of these parallel pathways and reveal mechanistic features relevant to the prospect of SIK inhibitor-based therapy.

## Results

### CREB1 and its paralogs drive cAMP-stimulated transcription

Signaling control of transcription downstream of cAMP-PKA activation occurs at the CREB1 paralogs, where PKA-phosphorylation and CRTC binding converge. To mechanistically examine this and create a testbed for resolving the relative contributions of CREB1 serine 133 phosphorylation and CRTC translocation, we generated a *Creb1*/*Atf1*/*Crem* triple knockout (hereafter "Creb1 TKO") line. This line lacks the full-length isoforms of these three paralogs, which together encode highly similar proteins featuring PKA-phosphorylated KID domains and bZIP domains that bind both CRTC coactivators and cAMP-responsive element (CRE) DNA sequences^2^.

Transcription profiling of Creb1 TKO cells with forskolin, an adenylyl cyclase activator that induces cAMP production, revealed that many strongly forskolin-responsive loci had substantially attenuated induction in Creb1 TKO cells compared to wild-type (71 out of 80 forskolin-responsive genes, |LFC|>2, *padj* < 0.01, Figure 1B and Supplemental Table 1). There were striking exceptions to this attenuation (*Hs3st1, Cdh6, Mitf)*, suggesting a CREB1 paralog-independent mechanism functions at some loci. Among these, *Mitf* was particularly notable, as it encodes a lineage-defining transcription factor for pigment cells that has been intensely studied as a direct CREB1 target gene (Figure 1A and Supplemental Figure 1A)^3,4^.

### CREB1 serine 133 phosphorylation is not required for CREB1 activity at endogenous loci

Having established that CREB1 paralogs are broadly required for forskolin-activated transcription, we next tested whether CREB1 serine 133 phosphorylation was necessary for this activity. We re-expressed human wild-type CREB1 or a phosphorylation-deficient CREB1^S133A^ mutant in Creb1 TKO cells (Figure 1C). Although a transfected CRE reporter showed reduced activity with CREB1^S133A^ compared to wild-type CREB1 (Figure 1D), both constructs rescued forskolin-induced transcription of the endogenous model locus *Sik1*, a well-documented target of cAMP-induced gene transcription^27^ to the same degree (Figure 1E-G and Supplemental Figure 1B). This suggests CREB1 phosphorylation by PKA is dispensable for CREB1 transcriptional activity; however, we reasoned this result could have been a specific feature of this target gene (*Sik1*), the timepoint (3 hr), or the high dose of forskolin used (10 µM). We extended our experiments comparing CREB1 and CREB1^S133A^ re-expressing cells across a panel of forskolin-responsive genes (*Mitf, Sik1,* and *Fos*) and a range of timepoints and doses of forskolin. In these experiments, we detected no consistent differences between CREB1 and CREB1^S133A^ across any condition tested (Figure 1F-G). Together, these results indicate that CREB1 serine 133 phosphorylation is dispensable for a high fraction of CREB1-dependent induction of endogenous target genes across a broad range of conditions.

### CRTC paralogs drive the cAMP-regulated transcriptome

Our findings regarding the dispensability of CREB1 phosphorylation pointed to CRTC as the alternative key transcriptional input downstream of cAMP-PKA. CRTC, like CREB1, has three paralogs: *CRTC1, CRTC2*, and *CRTC3* (*Crtc1*, *Crtc2*, and *Crtc3* in mice). To determine which CRTC paralogs mediated cAMP-PKA transcriptional responses, we leveraged the well-documented growth-suppressive effect of forskolin in melanoma (Figure 2A)^28^ and deployed a transcription-focused CRISPR-Cas9 library in human melanoma UACC257 cells to map genes required for this response (Figure 2B and Supplemental Table 2). In this screen, loss of *CRTC2* and *CRTC3* rescued growth while *CRTC1* did not detectably contribute (Figure 2B and Supplemental Table 2). None of the CREB1 paralogs scored, consistent with their redundancy described above. Across DepMap, *CRTC2* and *CRTC3* form a functionally distinct module from *CRTC1*, an effect most pronounced in melanoma but also detectable across other lineages (Supplemental Figure 2A-B)^29,30^. Guided by this understanding, we ablated *Crtc3* alone in B16F10 melanoma cells and saw partial loss of forskolin-induced *Mitf* transcription, which was completely abolished by co-ablation of *Crtc3* and *Crtc2* (Figure 2C and Supplemental Figure 2C). We observed similar results for the model target *NR4A2* in HEK293T cells upon co-ablation of *CRTC2* and *CRTC3* (Figure 2D and Supplemental Figure 2D).

**Figure 2.**
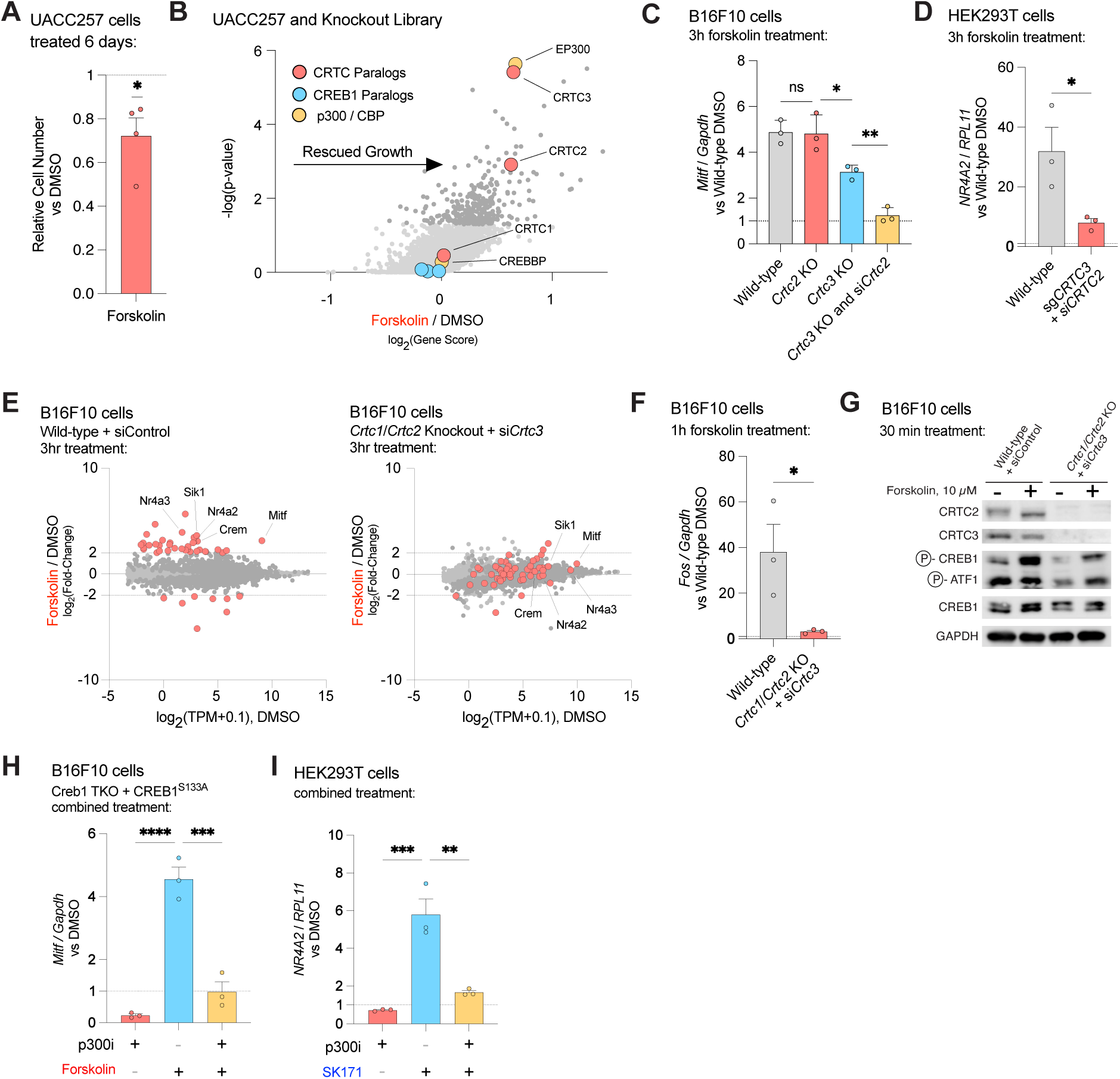
Loss of CRTC proteins disrupts cAMP-regulated transcription independently of CREB1 serine 133 phosphorylation. **(A)** Relative UACC257 cell number after 6 days of treatment demonstrating a mild cytostatic effect of forskolin (10 µM) versus DMSO (n = 3). **(B)** MAGeCK gene scores and nominal p-values from positive-selection CRISPR screen^73^. Dark gray points indicate *p* < 0.05. Genes whose loss rescued forskolin-inhibited growth (presumably through attenuated signaling) were enriched. CRTC, CREB1, and p300 paralog encoding genes are highlighted. **(C)** *Mitf* induction measured by RT-qPCR in wild-type B16F10 cells, single *Crtc2* and *Crtc3* gene knockout clones, and combined *Crtc3* knockout and *Crtc2* knockdown. Cells were transfected with indicated si*Crtc2* or non-targeting control siRNA as indicated. Samples were normalized to *Gapdh* and DMSO-treated cells; baseline levels were unchanged (n = 3, Supplemental Figure 2C). **(D)** *NR4A2* induction measured by RT-qPCR in wild-type HEK293T cells and combined bulk knockout/knockdown of *CRTC2* and *CRTC3.* Cells were transfected with indicated si*CRTC2* or non-targeting control siRNA as indicated. Samples were normalized to *RPL11* and wild-type DMSO-treated cells; baseline levels were unchanged (n = 3, Supplemental Figure 2D). **(E)** MA plots of transcriptional profiles from wild-type and *Crtc1/Crtc2* combined knockout + *Crtc3* knockdown B16F10 cells stimulated with 10 µM forskolin for 3 hr. Dark gray points indicate *padj* < 0.01. Red points highlight genes strongly regulated in wild-type cells (|LFC| > 2, *padj* < 0.01) and track their response upon compound CRTC paralog loss. Analysis was restricted to protein-coding mRNAs, fold-change and adjusted p-values were calculated using DESeq2. **(F)** *Fos* induction measured by RT-qPCR in B16F10 following a short, 1 hr forskolin (10 µM) treatment. Cells not transfected with si*Crtc3* were transfected with non-targeting control siRNA. Samples were normalized to *Gapdh* and DMSO-treated wild-type cells; baseline levels were unchanged (n =3, Supplemental Figure 2F). **(G)** Immunoblot of wild-type and *Crtc1/Crtc2* knockout + *Crtc3* knockdown B16F10 cells stimulated for 30 min with DMSO or forskolin (10 µM). **(H-I)** *Mitf* or *NR4A2* induction measured by RT-qPCR from B16F10 Creb1 TKO + CREB1^S133A^ cells or HEK293T cells, respectively. Cells were pretreated for 1 hr with the p300/CBP inhibitor A-485 (1 µM) before a 3 hr treatment with DMSO, forskolin (10 µM), or SK171 (1 µM). Samples were normalized to *Gapdh* or *RPL11* and DMSO-treated cells (n =3). *p<0.05, **p<0.01, ***p<0.001, ****p<0.0001. Bar plot statistical tests between conditions were assessed by one-way ANOVA with Sidak’s multiple comparison test, while comparisons to theoretical mean (i.e., 1.0) was performed by one-sample Students t-test. Error bars are standard error of the mean.

To test whether CRTC was required for the broader cAMP-regulated transcriptome, we generated *Crtc1/Crtc2/Crtc3* triple-ablated B16F10 cells. These cells largely phenocopied Creb1 TKO cells, showing attenuated activation of 63 out of 70 forskolin-responsive genes (|LFC|>2, *padj* < 0.01, Figure 2E, Supplemental Figure 2E, and Supplemental Table 3). Notably, *Mitf* and other genes that did not require the CREB1 paralogs for forskolin-responsive induction did require CRTC for their activation. (Figure 1B and 2E). This CRTC dependence held at early timepoints as well: *Fos* transcription after 1 hour of stimulation was abolished with CRTC loss (Figure 2F and Supplemental Figure 2F). Critically, forskolin-induced phosphorylation of CREB1 serine 133 remained intact in these cells (Figure 2G), demonstrating that phosphorylated CREB1 serine 133, the canonical mark of cAMP-induced gene activation, is insufficient to drive target gene transcription in the absence of CRTC.

CREB1 serine 133 phosphorylation recruits the paralogous histone acetyltransferases p300 and CBP through a direct physical interaction^8,10,11^. Our screen prominently identified *EP300*, which encodes p300 (Figure 2B), prompting us to ask whether p300 activity remained essential for PKA-CREB1-mediated transcription in the absence of CREB1 serine 133 phosphorylation. Chemical p300/CBP inhibition with A-485 blocked forskolin-driven *Mitf* transcription in Creb1 TKO cells re-expressing CREB1^S133A^, as well as SIK inhibitor-induced *NR4A2* activation in HEK293T cells (Figure 2H-I). Thus, p300/CBP remains essential for CREB1-driven transcription through recruitment mechanisms that are independent of serine 133 phosphorylation, likely through CRTC, as has been previously reported^31,32^.

### A positive, feedforward role for the annotated repressor ICER

Our CRTC-dominant model predicted that CREB1 bZIP domains are sufficient to recruit CRTC to DNA and activate transcription independent of CREB1 phosphorylation. A naturally occurring test of this principle exists: PKA drives the transcription of a set of truncated *CREM* isoforms collectively called ICERs (Inducible cAMP Early Repressors, Figure 3A)^33,34^. Because ICERs lack transactivation domains but retain their DNA/CRTC-binding bZIP domains, they have been characterized as repressors of PKA-CREB1 transcription that may compete with full-length CREB1 paralogs for CRE occupancy. However, if ICERs bind CRTC and CRE element, they could likely effect CRTC-directed transcription.

**Figure 3.**
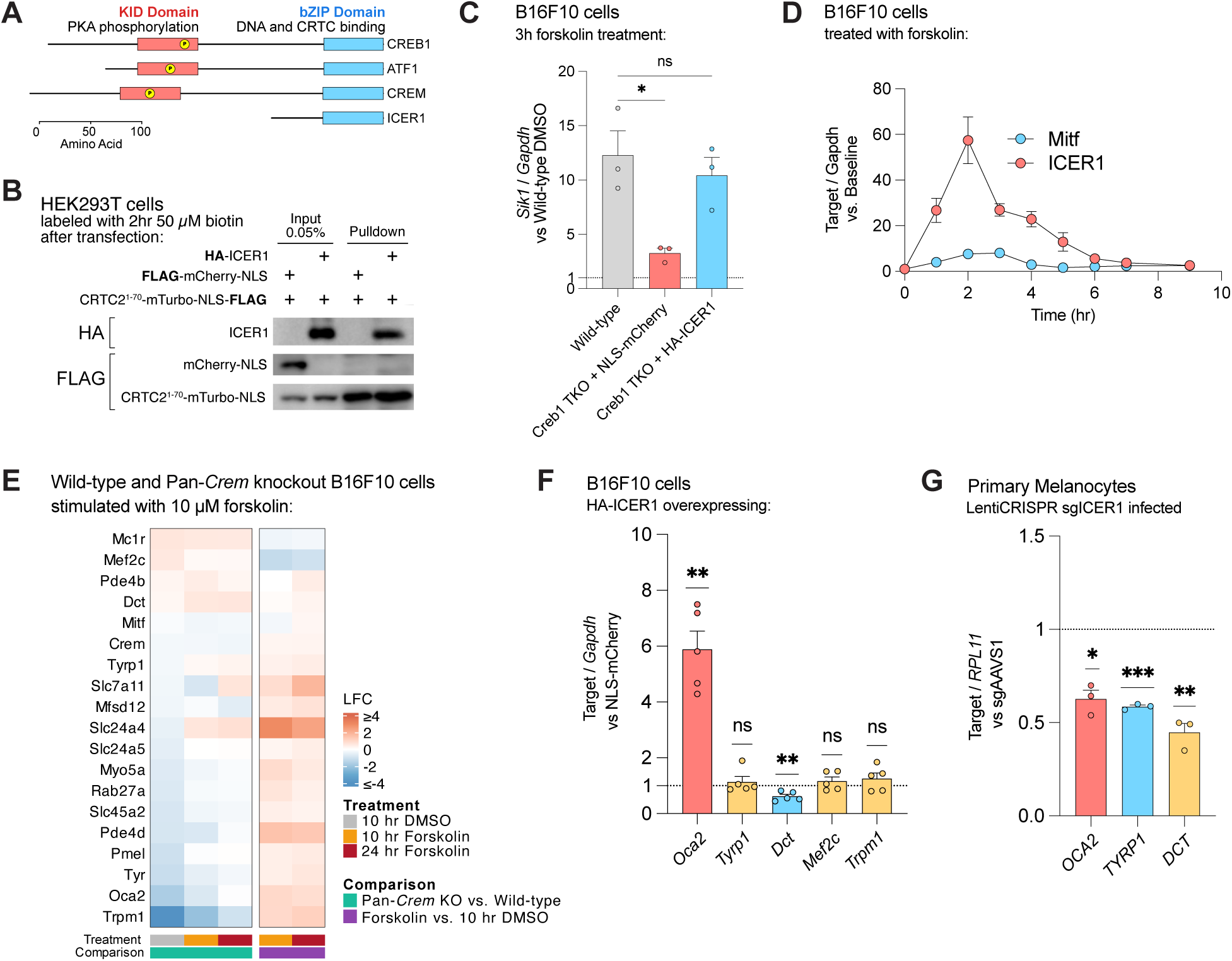
ICER1 positively regulates transcription and drives secondary pigment gene expression. **(A)** Domain diagrams of canonical CREB1, ATF1, and CREM compared to ICER1 (Uniprot: Q03060-8). **(B)** Proximity biotinylation of HA-ICER1 using CRTC2^1–70^-mTurbo-NLS fusion (which targets the CREB1-binding domain of CRTC2 to the nucleus) to assess interaction with HA-ICER1 versus a nuclear mCherry. FLAG-tagged mCherry-NLS and CRTC2^1-70^-mTurbo-NLS co-transfected constructs were resolved by differential gel migration. **(C)** *Sik1* induction measured by RT-qPCR in Creb1 TKO cells expressing HA-ICER1 or NLS-mCherry control, treated with DMSO or forskolin (10 µM). Samples were normalized to *Gapdh* and DMSO-treated wild-type cells; baseline levels were unchanged (n = 3, Supplemental Figure 2G). **(D)** Time-course analysis of ICER1 and *Mitf* induction in B16F10 cells upon forskolin treatment measured by RT-qPCR and normalized to *Gapdh* and the unstimulated baseline. ICER1 primers were designed against the first exon of the gene encoding the mouse ortholog of Uniprot: Q03060-8 (n = 3). **(E)** Heatmap of curated pigmentation genes in wild-type and pan*-Crem* knockout B16F10 cells at baseline (10 hr DMSO) and following forskolin stimulation (10 hr and 24 hr, 10 µM). **(F)** Pigmentation gene expression measured by RT-qPCR from B16F10 cells overexpressing HA-ICER1 versus NLS-mCherry control (n = 5). **(G)** Pigmentation gene expression measured by RT-qPCR in primary human melanocytes transduced with CRISPR-Cas9 targeting an ICER1-specific coding exon or a sgAAVS1 control. Cells were cultured in full melanocyte media (n = 3). *p<0.05, **p<0.01, ***p<0.001. Bar plot statistical tests between conditions were assessed by one-way ANOVA with Sidak’s multiple comparison test, while comparisons to theoretical mean (i.e., 1.0) was performed by one-sample Students t-test. Error bars are standard error of the mean.

We first tested whether ICER1 interacts with CRTC. A nuclear-localized miniTurbo (mTurbo) biotin ligase^35^ fused to the CREB1-binding domain of CRTC2 (residues 1-70) selectively labeled HA-ICER1 (ICER1, Q03060-8) but not a nuclear mCherry control (Figure 3B), consistent with ICER1 engaging CRTC2 through its bZIP domain. We then asked whether this interaction was sufficient for transcriptional activation and observed that re-expression of ICER1 in Creb1 TKO cells rescued forskolin-induced *Sik1* transcription to wild-type levels, phenocopying full-length CREB1 addback (Figure 3C and Supplemental Figure 2G).

In B16F10 cells, ICER transcripts (detected via an ICER1-specific first exon) are upregulated ∼60-fold in response to forskolin (Figure 3D), suggesting that ICER accumulation following an initial wave of cAMP-PKA activation could reinforce CRTC-dependent transcription as a feedforward amplifier. Because ICER proteins arise from multiple transcriptional start sites and some alternative translational start sites contained within shared *CREM/Crem* exons, we took complementary approaches to testing ICERs’ potential endogenous roles.

Frameshift mutation of a common *Crem* isoform exon in B16F10 cells, which retained intact *Creb1* and *Atf1*, caused a broad baseline reduction in genes activated by long-term cAMP activation including *Oca2*, *Trpm1,* and *Tyr* (Figure 3E, Supplemental Table 4), and 24 hr of forskolin treatment largely normalized these differences (Figure 3E, Supplemental Table 4). *Oca2* was the second most severely affected pigmentation gene, consistent with its previously reported unique sensitivity to *Crtc3* loss and with its strong reduction in our Creb1 TKO and CRTC triple-ablated experiments (Supplemental Figure 1A and 2E). Conversely, HA-ICER1 overexpression produced a ∼6-fold increase in *Oca2* but did not significantly increase other pigmentation genes assayed (Figure 3F). In contrast to most other pigmentation genes, *Dct* was elevated at baseline in *Crem-*mutant cells and repressed to ∼60% of control levels by HA-ICER1 overexpression (Figure 3F), suggesting that ICER1 still can function as a repressor depending on the locus or context. However, CRISPR targeting of ICER coding sequence in human primary melanocytes concordantly reduced all three pigmentation genes assayed: *OCA2, TYRP1, and DCT* (Figure 3G). Together, these loss- and gain-of-function results establish ICER1 as a net positive regulator of the broader cAMP-response pigmentation program.

### A transcriptomic survey of gene expression downstream of PKA and SIK-CRTC

We next sought to broaden the analysis of CRTC’s role in cAMP-induced transcriptional output beyond melanoma and HEK293T cells. We performed a transcriptomic survey across nine cell lines, comparing upstream cAMP-PKA activation to direct CRTC activation via pharmacological SIK inhibition. For SIK inhibition we used SK171 or HG-9-91-01^19,20^, and for upstream cAMP activation, since each cell type would require a different GPCR activating ligand, we used forskolin (Figure 1A)^36^. Our surveyed panel of lines included models of five lineages including melanocytes (primary human melanocytes, B16F10), hepatocytes (primary rat hepatocytes, HepG2), macrophages (PMA-differentiated THP1 cells), osteocytes (differentiated Ocy454 cells), and cortical neurons (primary mouse neurons). This diversity allowed us to detect common or overlapping signatures and to control for specialized cell culture conditions or transformation-associated signaling changes.

### Universal and cell type specific cAMP-regulated signatures

Cross-comparison of forskolin-stimulated cell models revealed a core set of CREB1-regulated genes shared across cell types. A total of 38 genes were concordantly upregulated and 7 were concordantly downregulated in at least 5 of 9 cell lines (|LFC| > 1, *padj* < 0.01; Figure 4A and Supplemental Tables 5-7). The most consistently upregulated genes were NR4A3 and *SIK1* (9/9), followed by *NR4A2*, *CREM*, *PDE4D*, and *CEBPB* (8/9). Motif enrichment analysis of the 38 universally upregulated genes identified full and partial CRE binding motifs as the dominant regulatory signature, with 12 of the top 15 enriched motifs corresponding to CRE-binding factors (Supplemental Figure 3A). Many of these genes often serve as model CREB1 targets^2^; collectively, we designate this 45-gene set the Universal cAMP-Activated Response (UCAR), and propose its use as a common reference signature for cross-lineage cAMP-PKA transcriptional outputs (Supplemental Table 7). This UCAR signature contains several paralogous and functionally overlapping groups of genes (Supplemental Figure 3B), including orphan nuclear receptors (*NR4A1*, *NR4A2*, *NR4A3*), dual-specificity phosphatases (*DUSP1*, *DUSP4*, *DUSP5*), cAMP phosphodiesterases (*PDE4B*, *PDE4D*, *PDE10A, PDE3A*), and bZIP transcription factors (*CREM*, *CEBPB*, *FOSL2*, *FOS*, *MAFF*, *JUNB*). Additionally, 9/45 (20%) of UCAR genes were identified recently as PKA inhibitor sensitive in PKA-driven cancers (32-fold enrichment, hypergeometric *p* = 7.4 x 10^-12^, Supplemental Table 7)^21^. Fewer genes were consistently downregulated by forskolin in the UCAR signature (*RNF144B, ARID5B, DEPP1, TXNIP, ZNF608, GPR37,* and *RGS3*) and no clear regulatory patterns were apparent for downregulation. This suggests that acutely after cAMP-PKA activation (3 hr), transcriptional activation likely dominates repressive signals.

**Figure 4.**
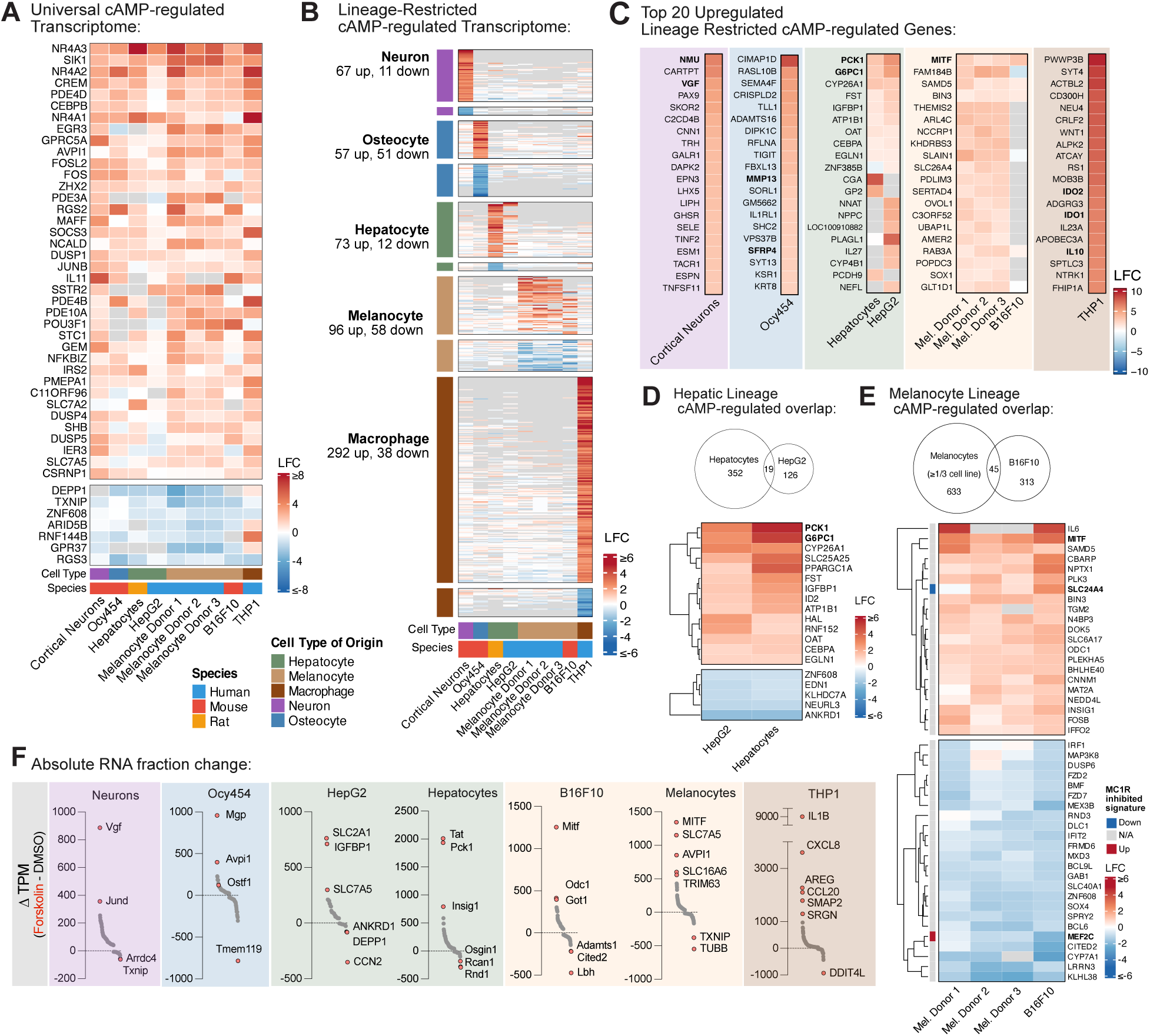
The cAMP-PKA axis activates universal and cell-type-specific gene expression signatures. **(A)** Heatmap of forskolin-induced differential gene expression after 3 hr of treatment. Genes that showed a consistent concordant change ≥5 cell lines are included (|log_2_FC| > 1, *padj* < 0.01). Rows are ordered by the number of cell lines in which each gene was differentially regulated (e.g., *NR4A3* and *SIK1* are upregulated in 9 of 9 lines). Gene labels denote human orthologs. **(B)** Heatmap of lineage-restricted cAMP-regulated transcriptomes (|LFC| > 2, *padj* < 0.01 in at least one cell line per lineage; not significant in other lineages). **(C)** Top 20 upregulated genes from (B). Well-documented lineage-defining factors discussed in the text are bolded. **(D–E)** Overlap of concordantly regulated genes between primary hepatocytes and HepG2 cells or primary melanocytes and B16F10 cells (|LFC| > 1, *padj* < 0.01). In the melanocyte comparison, the leftmost column indicates each gene’s status in the previously described *Mc1r-*inhibited melanocyte gene signature^47^. **(F)** Absolute change in RNA abundance (ΔTPM). The top and bottom quartiles of ΔTPM values are shown for each lineage’s significantly regulated genes (|LFC| > 1, *padj* < 0.01). Genes with the largest ΔTPM values or notable biological significance are labeled. All comparisons in (A-F) derive from *n* = 3 biological replicates per condition. Forskolin (10 µM) was compared to matched DMSO vehicle controls. Analyses were restricted to annotated protein-coding genes. Fold-changes and Benjamini-Hochberg–adjusted *p*-values were generated by DESeq2, run independently for each cell line and treatment. TPM values were calculated from raw counts for abundance estimation.

Lineage-restricted gene expression changes were substantially more diverse than the UCAR signature. Hundreds of genes were differentially regulated exclusively within each of the 5 lineages with a 2-fold cutoff and dozens to hundreds with a 4-fold cutoff (*padj* < 0.01; Figure 4B and Supplemental Figure 3C). This increased breadth we observed when compared to UCAR was not simply a statistical artifact of comparing fewer cell lines per lineage: among three melanocyte cell preparations from independent donors, 115 non-UCAR genes were concordantly regulated in all three donors (|LFC| >1, *padj* <0.01). These programs included canonical lineage-defining targets: melanocytes upregulated the pro-pigmentation gene *MITF*, THP1-derived macrophages induced the anti-inflammatory gene *IL10*, and hepatocytes upregulated the gluconeogenic genes *G6PC1* and *PCK1* (Figure 4C)^1,2,5,8,37–39^. Neurons upregulated brain-restricted transcripts encoding neuronal signaling molecules including *Nmu* (neuromedin U) and *Vgf* (VGF nerve growth factor)^40,41^, while differentiated Ocy454 osteocytes showed a strong increase in the bone-specific matrix metalloprotease *Mmp13* and *Sfrp4* (Figure 4C)^42,43^.

Transformed cell lines and primary counterparts derived from a common lineage (HepG2 and hepatocytes; B16F10 and melanocytes) had surprisingly low overlap in their lineage-restricted responses (∼10% for both lineages, Figure 4D-E). This weak overlap most likely reflects signaling alterations that occur with transformation or potentially differences in media conditions like those required for primary cell culture. Nonetheless, the intersection of tumor and primary cell responses contained well-documented model targets and appeared to represent the most robust "hard-wired" cAMP responses that exists for each lineage. On the other hand, primary cells had richer responses overall, with twice as many genes modulated by forskolin, many with cell type relevance. In melanocytes, these included factors known to be involved in pigmentation including *ADRB2, SLAIN1,* and *WIPI1*^44–46^, while hepatocytes uniquely upregulated metabolic enzymes and signaling kinases that support gluconeogenesis through amino acid breakdown, including *Tat, Got1*, and *Ppm1k*, which were not substantially changed in HepG2 cells.

The survey also revealed that lineages differ not just in which genes are regulated but in the transcriptional architecture through which cAMP signaling engages effector programs. Hepatocytes and THP1 macrophages directly upregulated effector genes such as gluconeogenic enzymes (*PCK1*, *G6PC1*) and *PPARGC1A* in the case of hepatocytes or cytokines (*IL10*) and phagosome/lysosome (*RAB20, LAMP3*) genes in the case of macrophages. Melanocytes, in contrast, acutely modulated only 2 genes directly associated with long-term cAMP-related pigmentation changes^47^. Instead, this lineage relies on *MITF* to drive a secondary wave of melanosome biogenesis and melanin biosynthesis gene transcription (Figure 4E).

We reasoned there might also be meaningful changes in the expression of highly expressed genes that were not apparent in our ratiometric analysis. Examining changes in total transcript abundance (ΔTPM) revealed several such cases across lineages (Figure 4F). For instance, in osteocytes, the bone-development genes *Mgp* and *Ostf1* showed modest fold-changes (∼3x) but substantial absolute shifts. *SLC7A5* and *SLC16A6* were upregulated ∼500-1000 TPM units in melanocytes. These genes represent plasma membrane and melanosomal tyrosine uptake systems, respectively, which likely enable melanin synthesis in response to cAMP activation^48,49^. One of the strongest changes we identified was the upregulation of pro-inflammatory *IL1B* and *CXCL8*, which increased by several thousand TPM units in THP1-derived macrophage cells alongside the much stronger ratiometric changes of immunosuppressive transcripts like *IL10* and *IDO1*^50^. THP1-derived macrophages are an imperfect model of primary macrophages, but the potential co-regulation of opposing inflammatory programs by cAMP-PKA warrants further attention^22^.

### Genome-wide concordance between forskolin and SIK inhibition across cell types

Having defined the shared and lineage-specific transcriptional outputs of cAMP signaling, we next asked how faithfully chemical SIK inhibition could recapitulate this response. Across cell lines treated with SIK inhibitors in parallel, we found that forskolin and SIK inhibition gene expression programs correlated highly, with lineage identity, rather than treatment, driving most of the variation between conditions (Figure 5A-B). Profiles induced by HG-9-91-01 and SK171 near-perfectly correlated within the same cell line (slope = 0.94, r = 0.99, Supplemental Figure 4A), suggesting these drugs are functionally interchangeable. We found no evidence for discrete subclasses of forskolin- and SIK inhibitor-responsive genes; both UCAR and cell type-restricted transcripts were induced by SIK inhibition, and no identifiable gene class showed preferential responsiveness to either treatment (Figure 5B and Supplemental Figure 4B). Consistent with the highly correlated expression profiles, genes strongly induced by forskolin (>4-fold) were also differentially expressed with SIK inhibition (∼60-100% of loci, Figure 5C). Neurons were a notable exception, having a severely blunted response to SIK inhibition and a correspondingly reduced overlap with the forskolin program (Figure 5C and D). To test whether our readout could resolve overlapping but distinct transcriptional programs, we compared forskolin stimulation to KCl-mediated depolarization in cortical neurons and found while they produced correlated responses, many genes showed an asymmetric preference for one stimulus over the other contrasted to forskolin versus SIK inhibitor (Supplemental Figure 4B–C).

**Figure 5.**
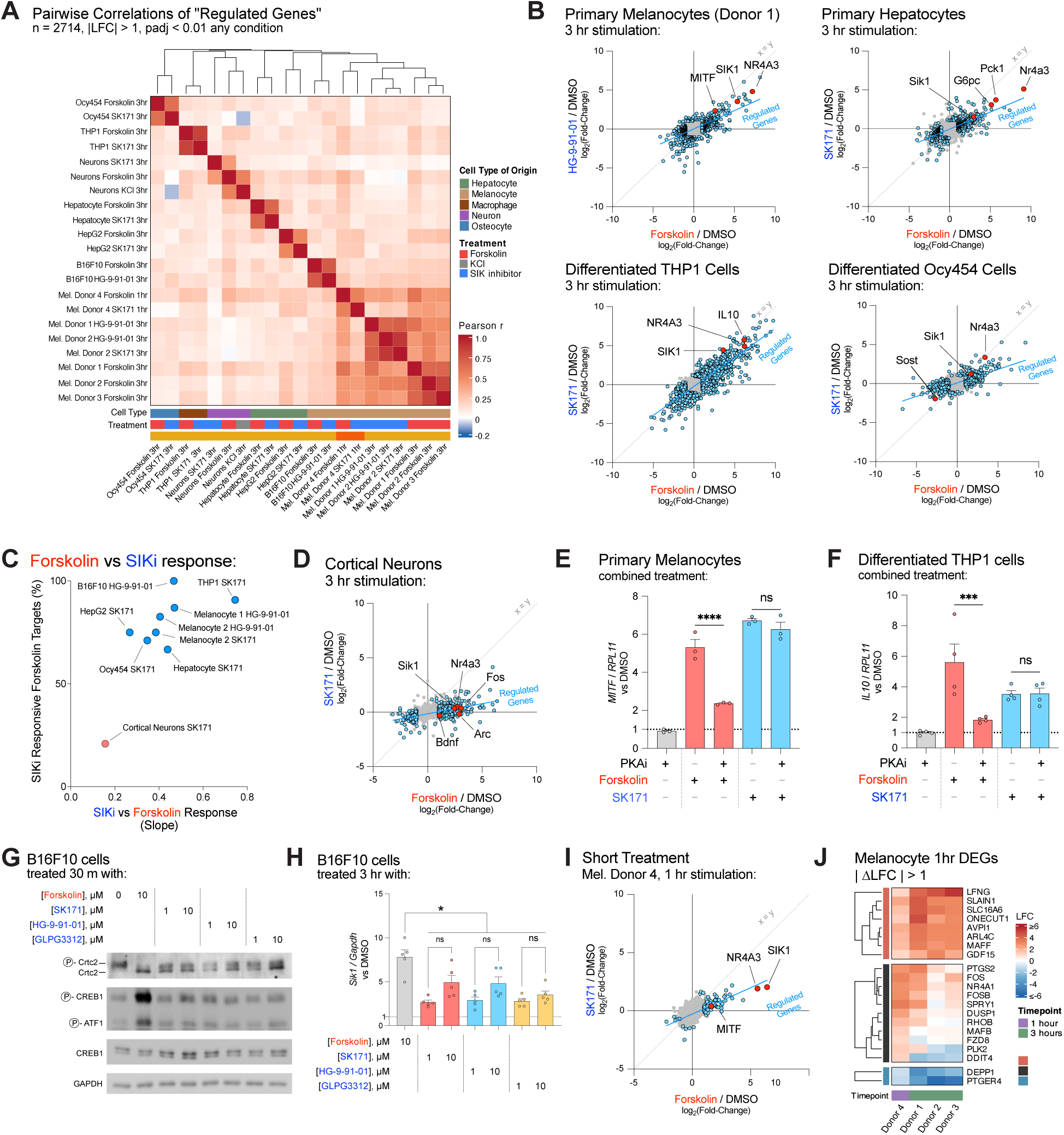
Pharmacological SIK inhibition largely recapitulates upstream cAMP-induced transcription. **(A)** Pearson correlation matrix of all cell lines and treatments focused on the 2714 genes (and their orthologs) that were differentially expressed in at least one sample (|LFC|>1, *padj* <0.01). **(B)** Representative scatterplots comparing transcriptional responses to forskolin (10 µM) versus SIK inhibitor treatment across profiled cell lines after 3 hr. Blue points indicate differentially expressed genes under either treatment (|LFC| > 1, *padj* < 0.01). The blue line represents a linear fit to these points. Red points highlight select universally regulated genes *SIK1* and *NR4A3* (and their orthologs) and lineage-defining PKA-controlled genes. **(C)** Comparison of the SIKi-versus-forskolin regression slopes (*x*-axis) and the percentage of strongly forskolin-responsive genes (|LFC| > 2, *padj* < 0.01) that are also significantly SIKi-responsive at any magnitude (TPM > 1 baseline filtered, *padj* < 0.01). **(D)** Scatterplot of genes from cortical neurons treated with SK171 (1 µM) or forskolin (10 µM). Points and linear fit as in (B). **(E)** *MITF* induction measured by RT-qPCR from primary melanocytes. Cells were pretreated for 1 hr with the PKA inhibitor BLU0588 (2 µM) before a 3 hr treatment with forskolin (10 µM) or SK171 (1 µM). Samples were normalized to *RPL11* and DMSO-treated cells (n =3). **(F)** *IL10* induction measured by RT-qPCR in THP1 derived macrophages, treated as in (E) (n = 4). **(G)** Immunoblot of B16F10 cells treated for 30 min with forskolin (10 µM) or the SIK inhibitors SK171, HG-9-91-01, or GLPG3312 (1 µM and 10 µM). **(H)** *Sik1* induction measured by RT-qPCR from B16F10 cells, normalized to *Gapdh* and DMSO control (n = 5). **(I)** Scatterplot showing correspondence between forskolin (10 µM) and SIK inhibitor treatment after 1 hr in primary melanocytes. Points and linear fit as in (B). **(J)** Heatmap of genes that were differentially expressed after 1 hr with |LFC| > 1 compared to the average LFC after 3 hr in melanocytes. All RNAseq analyses were restricted to annotated protein-coding genes. Fold-changes and Benjamini-Hochberg–adjusted *p*-values were generated by DESeq2, run independently for each cell line and treatment. *p<0.05, **p<0.01, ***p<0.001, ****p<0.0001. Bar plot statistical tests were from ordinary, one-way ANOVA with Sidak’s multiple comparison test. Error bars are standard error of the mean.

SIK inhibitor-induced transcription activation was uniformly weaker than forskolin treatments across all cell lines, as reflected by regression slopes consistently below 1.0 (Figure 5B and 5D). This blunted transcriptional response to SIK inhibition compared to forskolin could reflect incomplete CRTC activation by SIK inhibition or a contribution from basal PKA-mediated CREB1 phosphorylation. Across time-points, cell lines, and loci, a pre-treatment with PKA inhibitor did not affect responses to SIK inhibition (Figure 5E-F and Supplemental Figure 4D-F), arguing against a required contribution from basal PKA-mediated CREB1 phosphorylation. Instead, consistent with incomplete suppression by SIK inhibitors, 30 min of SIK inhibitor treatment produced incomplete CRTC2 dephosphorylation, as assessed by gel-shift in B16F10 and HEK293T cells (Figure 5G and Supplemental Figure 4G). This held even when inhibitor concentrations were increased 10-fold or when cells were treated with the pan-SIK1/2/3 inhibitor GLPG3312 (HG-9-91-01 and SK171 are ∼10x selective for SIK2 and SIK3 over SIK1).^19,20^ Transcriptional induction of CREB1 targets (*Sik1* in B16F10, *NR4A2* in HEK293T) likewise remained submaximal across all conditions tested (Figure 5H and Supplemental Figure 4H). The attenuated transcriptional response to SIK inhibition likely reflects a difference in pharmacology (perhaps incomplete or slower target engagement kinetics with SIK inhibitor treatment) rather than a missing contribution from CREB1 phosphorylation.

Mirroring our genetic analysis, we also asked whether a shorter treatment duration might reveal separate forskolin and SIK inhibitor responses. Although a somewhat different set of genes was upregulated at 1 hr than at 3 hr, the SIK inhibitor and forskolin responses again mirrored one another at each timepoint, revealing no separation even at the earlier timepoint (Figure 5I-J). Additionally, the degree of SIK inhibitor response was similar at both timepoints (Slope of 0.39 versus an average of 0.42, Figure 5B and 5J). Approximately 7-fold fewer genes were differentially expressed at the earlier timepoint. Among genes showing notable differences between 1 hr and 3 hr treatments, we identified two patterns (albeit from a non-isogenic comparison): 1) genes not yet at peak expression at 1 hr, 5/8 of which corresponded to genes with large absolute changes in our ΔTPM analysis (Figure 5I-J and Figure 4F), and 2) genes more strongly induced at 1 hr than at 3 hr, likely reflecting early negative feedback. The immediate early gene *FOS* exemplifies the latter, as it also returns to near baseline after 3 hr of activation in B16F10 cells (Figure 5I-J and Figure 1F).

## Discussion

Here we demonstrate that the SIK-CRTC axis, not CREB1 phosphorylation, provides the dominant input for CREB1 transcriptional activity across many cell types. We propose that SIK-CRTC should therefore be considered the canonical activation model of PKA-CREB1-controlled transcription, displacing PKA-mediated CREB1 phosphorylation from that role. A non-phosphorylatable CREB1 mutant, in cells lacking the redundant CREB1 paralogs ATF1 and CREM, did not detectably alter transcriptional responses to forskolin. Conversely, phosphorylated CREB1, in the absence of CRTC proteins, was unable to activate endogenous gene expression in response to PKA activation. When overexpressed, ICER, a *CREM* isoform that lacks the PKA-phosphorylation site, could substitute for the full-length CREB1 paralogs and drive cAMP-PKA-regulated gene expression through CRTC. Finally, as a pharmacological test of our genetic findings, chemical SIK inhibition activated the majority of cAMP-PKA-induced gene expression across diverse cell types, suggesting these results generalize beyond any single lineage. As part of this survey, we also provide a cross-cell-type catalog of PKA-induced transcriptional responses spanning melanocyte, hepatocyte, macrophage, osteocyte, and neuronal lineages, which reveal a small core group of genes regulated by cAMP across lineages alongside more substantial lineage-specific gene sets.

CREB1 serine 133 conservation extends at least to cnidarians (as does CRTC)^12^, which argues for its functional importance, as do decades of research into phosphorylated CREB1 function. Given this, why does it appear to be dispensable in numerous lineages for cAMP-PKA activated transcription? First, much of the foundational evidence establishing CREB1 phosphorylation’s importance comes from studies performed in the nervous system, where CaMK-dependent phosphorylation plays a larger role^51,52^. Our findings that SIK inhibition has minimal transcriptional activity in cortical neurons are consistent with the divergent CREB1 regulation in these cells. This is further supported by the documented preferential role of CRTC1 over CRTC2 and CRTC3 as a key input for neuronal function, suggesting CRTC paralog specialization with neuronal identity as a dividing line ^53,54^. Other data suggest that additional cell lineages or cells from developmentally distinct cell states might also preferentially require CREB1 phosphorylation^55^.

Beyond cell type differences, certain methods and tools might have also complicated our understanding of CREB1 phosphorylation. CREB1^S133A^ has been used as a dominant negative, and while it may behave differently from wild-type CREB1 at high expression levels^2,56–58^, our findings suggest that CREB1^S133A^ is transcriptionally active, consistent with related reports^59^. Another widely used tool has been transfected CRE reporter plasmids, which we found do detect a difference between wild-type and CREB1^S133A^. These reporters are present at high copy numbers, are supercoiled, and lack full genomic context. These nonphysiological properties might explain why this tool led to overestimation of the activating signal from phosphorylated CREB1.

Finally, the known molecular function of CREB1 phosphorylation must be partially masked by parallel redundant mechanisms. While the interaction of p300 and phosphorylated CREB1 has been characterized in atomic detail, our findings suggest p300/CBP does not require phosphorylated CREB1 for its action in PKA-mediated transcription. This is consistent with other reports of a cooperative p300 recruitment mechanism that acts through CRTC^31^. More broadly, conservation of a phosphorylation site does not necessitate a dominant regulatory role. This parallels rpS6 phosphorylation, which is conserved back to yeast yet produces only mild phenotypes when ablated in mice^60^. PKA-mediated CREB1 phosphorylation may play a similar modulatory role.

ICER1, a naturally occurring *CREM* isoform lacking the PKA phosphorylation site^34^, provided a minimal test of whether CRTC recruitment alone is sufficient for transcriptional activation. Our experiments suggest that ICER1 drives CRTC-mediated transcription. This finding contrasts with the previous description of ICER as a transcriptional repressor^34,61^; however, the two mechanisms are not mutually exclusive: ICER may activate transcription at moderate levels through CRTC recruitment and shift to a squelching inhibitor at high expression levels where CRTC is limiting. Together with the autoregulation of ICERs by PKA-CREB1, this dual capacity warrants further investigation of the contexts where ICER acts as a positive versus negative regulator. More broadly, this finding is a reminder that isoform-level complexity at loci like *Crem* can elude gene-focused tools such as pre-designed CRISPR and siRNA reagents.

Our findings suggest that in many cell types, chemical SIK inhibition can recapitulate much of the transcriptional output of GPCR activation, without a significant transcriptional bias toward particular target genes. Chemical SIK inhibition activated gene expression somewhat less robustly as compared to forskolin, and this ceiling was not substantially raised by increasing inhibitor concentrations or by using the pan-SIK1/2/3 inhibitor GLPG3312, suggesting this limitation is a fundamental limitation of chemical SIK inhibition. Non-transcriptional effects elicited by forskolin downstream of cAMP-PKA might contribute to these differences in activation strength and kinetics^62,63^. While this complicates direct comparison with forskolin, it does not represent a limitation for SIK inhibitor therapy. Chemical SIK inhibition is a more direct approach to activate CRTC-dependent gene expression than receptor- or adenylyl cyclase-level pathway activation, which is complicated by the diverse non-transcriptional consequences and targets of global PKA activation^64^ and cAMP signaling compartmentalization^65,66^. It remains to be seen whether alternative SIK targeting strategies such as degraders or allosteric modulators may activate the pathway with increased efficacy versus ATP-competitive inhibition. While the partial activation we observe may set an inherent efficacy ceiling for indications requiring full pathway engagement, it may equally provide a built-in safety margin against dose-limiting toxicities.

Our work systematically defines a mechanistic fact that has been emerging from work in multiple systems over decades: that SIK-CRTC is the dominant input for CREB1 transcriptional activation across many cell types, loci, and timepoints^1,2,8,21,37,39^. A simplified model of PKA-CREB1 activation cannot cover all the nuances that clearly exist between cell types and between the CRTC and CREB1 paralogs, but we propose that the default model for most cell types should emphasize the SIK-CRTC pathway over CREB1 serine 133 phosphorylation. This revision has concrete implications for SIK inhibitor development: the incomplete CRTC activation we observed defines a therapeutic ceiling that may need to be overcome for maximal efficacy and the cell type-specific transcriptional responses we describe can guide predictions of on-target toxicities across tissues.

## Acknowledgements

We thank Priya P. Budde, Stephen M. Ostrowski, Magdy Gohar, and Kiran Ebrahimi for critical feedback on the manuscript and Inbal Rachmin and Paul C. Rosen for their input during early stages of the project, as well as early experimental support from Christopher D. George. We are also thankful for input from the entire Fisher lab, especially Jessica L. Flesher, Sharon Germana, and Andrew King. We also thank the Broad Institute Clinical Research Sequencing Platform and the Saint John’s Cancer Institute Genomic Sequencing Center for sequencing services supported from the Dr. Miriam and Sheldon G. Adelson Medical Research Foundation (AMRF). C.H.A. was supported by the Damon Runyon Cancer Research Foundation (DRG-2454-22) and NIAMS (K99 AR084551). D.E.F. acknowledges support to his laboratory from NIH grants P01 CA163222, R01 AR072304, and R01 AR043369, as well as funding from the Dr. Miriam and Sheldon G. Adelson Medical Research Foundation, the Melanoma Research Alliance, the Lancer Professorship at Harvard Medical School, and the Water Cove Charitable Foundation. M.N.W. acknowledges support from NIDDK (R01DK116716) and NIAMS (P50AR080596) and a Chen Institute MGH Research Scholar award. C.S. was supported by the Helen Hay Whitney Foundation. G.P. acknowledges support from NIAMS (Clinical Investigator Award K08AR084618). D.S.B.H. and the Genomic Sequencing Center funding was from the Dr. Miriam and Sheldon G. Adelson Medical Research Foundation (AMRF).

## Author Contributions

Conceptualization, C.H.A. and D.E.F.; Investigation, C.H.A., M.L., A.E.G., L.Hu., J.R.B., A.V., X.W., E.Z., J.T., A.A., J.P.C., S.R., C.S., L.Hy., T.V., and G.P.; Formal Analysis, C.H.A.; Visualization, C.H.A.; Writing – Original Draft, C.H.A.; Writing – Review & Editing, C.H.A. and D.E.F.; Supervision, D.S.B.H., M.N.W., N.H., and D.E.F.; Funding Acquisition, C.H.A., D.S.B.H., M.N.W., N.H., and D.E.F.

## Methods

### Cell Culture

All cell culture was performed at 37°C and 5% CO_2_ except for pre-differentiated Ocy454, which were propagated under the permissive temperature of 33°C before differentiation.

**HEK293T cells** were the "Lenti-X" subclone obtained directly from Takara Bio (#632180). Cells were cultured in DMEM (Invitrogen, #11995073) supplemented with 10% v/v heat-inactivated fetal bovine serum (BioTechne, #S11550H) and 100 U/mL penicillin-streptomycin (Invitrogen, #15140163) and 50 µg/mL Normocin (Invivogen, #ant-nr-2). For all experiments, cells were treated in complete medium.

**B16F10 cells** were obtained from ATCC. Cells were cultured in DMEM (Invitrogen, #11995073) supplemented with 10% v/v heat-inactivated fetal bovine serum (BioTechne, #S11550H) and 100 U/mL penicillin-streptomycin (Invitrogen, #15140163) and 50 µg/mL Normocin (Invivogen, #ant-nr-2). For all experiments, cells were treated in complete medium.

**UACC257 cells** were obtained from the MGH Center for Molecular Therapeutics^67^. Cells were cultured in DMEM (Invitrogen, #11995073) supplemented with 10% v/v heat-inactivated fetal bovine serum (BioTechne, #S11550H) and 100 U/mL penicillin-streptomycin (Invitrogen, #15140163) and 50 µg/mL Normocin (Invivogen, #ant-nr-2).

**HepG2 cells** were obtained from the MGH Center for Molecular Therapeutics^67^. Cells were cultured in DMEM (Invitrogen, #11995073) supplemented with 10% v/v heat-inactivated fetal bovine serum (BioTechne, #S11550H) and 100 U/mL penicillin-streptomycin (Invitrogen, #15140163) and 50 µg/mL Normocin (Invivogen, #ant-nr-2). For all experiments, cells were treated in complete medium.

**THP1 cells** were obtained from the MGH Center for Molecular Therapeutics^67^. Cells were cultured in RPMI-1640 medium (Invitrogen, #11875119) supplemented with 10% v/v heat-inactivated fetal bovine serum (BioTechne, #S11550H), 100 U/mL penicillin-streptomycin (Invitrogen, #15140163), and 50 µg/mL Normocin (Invivogen, #ant-nr-2). THP1-derived macrophage-like cells were generated by treating cells (1 × 10^6^ per well in a 6-well plate) with 1 µg/mL PMA for 24 hr before PMA was removed the night before treatment.

**Ocy454 cells** were previously generated and cultured as described^68^. Cells were maintained at permissive temperature of 33°C and 5% CO_2_ in MEM (Invitrogen, # 11095080) supplemented with 10% fetal bovine serum and 1% antibiotic and antimycotic. For differentiation, confluent cells were shifted to 37°C for 7 days before treatment.

**Primary Wistar rat hepatocytes** were obtained freshly isolated from the Massachusetts General Hospital Cell, Tissue, and Organ Resource Center and were plated on rat-tail collagen coated 6-well plates overnight in complete hepatocyte medium, provided by the core and previously described^69^. The next evening, medium was exchanged to Williams’ E medium (Invitrogen, #12551032) supplemented with glutaMAX (Invitrogen, #35050061) and 100 U/mL penicillin-streptomycin (Invitrogen, #15140163). Cells were treated and harvested the following day.

**Primary human foreskin melanocytes** were isolated from neonatal foreskin according to institutional rules for anonymized discarded tissue acquisition. Epidermis was isolated by gentle peeling from foreskins after overnight digestion at 4°C with 3 parts 5 U/mL dispase solution (StemCell Technologies, #07913) and 1 part 0.25% trypsin-EDTA (Invitrogen, #25200114). Epidermis was then dissociated in 0.05% trypsin-EDTA solution at 37°C for 15 min before trituration with a 10 mL serological pipette and filtering through a 70 µm cell strainer. Cells were plated and expanded in M254 medium (Invitrogen, #M254500) supplemented with 100 U/mL penicillin-streptomycin (Invitrogen, #15140163) and 1× HMGS (Invitrogen, #S0025). After 7 days and several media changes, purity was evaluated by inspection of melanocyte morphology, and cultures with purity estimated to be >90% were moved forward for assay. Medium was exchanged to M254 without HMGS the night before all treatments (except sgICER harvest, which was performed in complete medium).

**Primary mouse cortical neurons** were isolated from FVB embryonic mice harvested at E16.5. Cortical tissue was dissected and dissociated and plated on Neuronal Coating Solution coated plates (Sigma Aldrich, #02705) in Neurobasal Media (Life Technologies #21103049) supplemented with 100 U/mL penicillin-streptomycin (Invitrogen, #15140163), 1x GlutaMAX (Invitrogen, #35050061), and 1x B-27 supplement (Invitrogen #17504044) with media exchange after 4 hours. After 7 days of post-isolation maturation, cells were silenced with overnight treatment with APV (100 µM) followed by 1 hr of lidocaine (300 µM) pretreatment before stimulation. For depolarization, 3x depolarization buffer was added to a final concentration of 55 mM KCl, 350 µM MgCl_2_, 700 µM CaCl_2_, and 3.5 mM HEPES, pH 7.4.

## Chemical Treatments

All chemicals were stored at 2,000× their final working concentration in DMSO and all final DMSO concentrations were 0.05% (v/v). SK171 was synthesized as previously described^19^. Forskolin was from Santa Cruz Biotechnology (#SC-3562A). HG-9-91-01 and PMA were from Sigma-Aldrich (#SML3029, #524400). A-485, BLU0588, and GLPG3312 were from MedChemExpress (#HY-107455, #HY-153967, and #HY-157442). Lidocaine and APV were from Selleck (#S1357 and #E2979).

For time-resolved experiments, treatments were applied in reverse chronological order so that all samples were harvested simultaneously.

## Knockout and Stable cDNA Expression

Knockout cell lines were generated using the pLentiCRISPRv2-Opti vector (Addgene #163126) with the guide sequences listed below. For clonal knockout generation, CRISPR plasmid was transfected into cells using Lipofectamine 3000 (Invitrogen, #L3000008). 48 hours after transfection, cells were treated with puromycin (0.5 µg/mL) for 48–72 hours and single-cell cloned via FACS. Knockout clones with frameshift indels were identified via next-generation amplicon sequencing.

For viral CRISPR-Cas9 knockout populations and addback expression experiments, lentivirus was generated in HEK293T cells by transfecting VSV-G, psPAX2, and transfer vector in a 1:3:3 ratio by weight in either a 6-well or 15 cm format. Virus was harvested 48 hours after transfection, frozen, and used for subsequent infections. For primary melanocyte infection, virus was first concentrated by adding 1 part viral supernatant to 3 parts viral concentrator solution (40% w/v PEG-8000, 1.2 M NaCl) and incubating overnight at 4°C, before centrifugation at 1,600 × *g* at 4°C for 60 min and resuspended in PBS.

Stable expression experiments utilized the lentiviral backbone pLJC5 containing GFP (Addgene #255533), human CREB1 (Addgene #254872), or CREB1^S133A^ (Addgene #254871) as well as the lentiviral backbone pTwist Lenti-SFFV-WPRE containing FLAG-mCherry-NLS (Addgene #254873), HA-ICER1 (Addgene #254874), and CRTC2^1–70–mTurbo–NLS–FLAG^ (Addgene #254875).

## CRISPR Screen

UACC257 cells were infected with a custom CRISPR-Cas9 knockout library cloned into pLentiCRISPRv1 that encoded 28,697 guides targeting 6,370 genes enriched for those encoding transcription factors, metabolic genes, and ubiquitin ligases and ubiquitin-ligase like pathways^70,71^. Cells were infected such that ∼1000 cells were infected per guide. After four days of selection in 1 µg/mL puromycin, cells were split into 2x treated arms: 0.05% (v/v) DMSO and 10 µM forskolin in an equivalent volume of DMSO. Treatments were refreshed upon passage (every 2-3 days). Both arms were harvested after 10 doublings. Next-generation sequencing libraries were amplified and sequenced as previously described^72^. Reads with exact guide matches were analyzed via MAGeCK (v0.5.9.5)^73^.

## siRNA-Mediated Knockdown

Four-plex "SMARTpool" siRNAs from Dharmacon were used for knockdown experiments. 5 × 10^5^ cells were plated and transfected with 20 pmol of siRNA using Lipofectamine RNAiMAX (Invitrogen, #13778150). Cells were split the next day into individual wells for treatment and harvested the following day (48 hours post-transfection) for western blotting or RT-qPCR.

## RNA Isolation and qPCR

For RT-qPCR experiments, 3 × 10^5^ cells were plated in 12-well plates, incubated overnight, and treated the next day. For RNAseq, 1 × 10^6^ cells were plated in 6-well plates, incubated overnight, and treated the next day. In both formats, cells were lysed in Qiagen RLT buffer and RNA was purified via Qiagen RNeasy columns (Qiagen #74106).

For RT-qPCR experiments, biological replicates were treated and harvested independently (i.e., on different days). After isolation, transcripts were quantified via one-step RT-qPCR kits from KAPA/Roche according to the manufacturer’s instructions. Fold-change values were calculated from the median C_T_ value of target and control genes (generated in technical triplicate for each biological replicate).

## RNA-seq

RNA was isolated according to the same protocol as above. All RNAseq replicates are from independent samples treated alongside each other (i.e., on the same day in parallel). Two separate preparation protocols were used in our RNA-seq survey across two institutions. RNA-seq libraries were prepared using one of two protocols depending on the sequencing site, as detailed in Supplemental Table 8.

*Protocol 1 (Genomic Sequencing Center, Saint John’s Cancer Institute).* Extracted RNA quality and quantity were assessed using the Agilent TapeStation 4200 System with High Sensitivity RNA ScreenTape (Agilent Technologies) and Qubit 4 Fluorometer (Invitrogen), respectively. Libraries were constructed using the Stranded mRNA Prep, Ligation kit (Illumina) following manufacturer’s recommendations. Sequencing libraries were assessed for quality and quantity using the Agilent TapeStation 4200 System with High Sensitivity D1000 ScreenTape (Agilent Technologies) and Qubit 4 Fluorometer (Invitrogen). Libraries were normalized to 4 nM, pooled, denatured, and loaded onto an Illumina NextSeq 550 or NextSeq 2000 at 1×76 bp (single-end) or 2×76 bp (paired-end) read specification. FASTQ files were generated from sequencing data using the Illumina BaseSpace Sequence Hub and DRAGEN Illumina software.

*Protocol 2 (Broad Institute).* RNA was quantified using the Quant-iT RiboGreen RNA Assay Kit (Invitrogen, #R11490) and normalized to 5 ng/µL. An aliquot of 325 ng from each sample was processed for library preparation using an automated variant of the Illumina TruSeq Stranded mRNA Sample Preparation Kit, which preserves strand orientation using oligo(dT) bead selection of mRNA from total RNA followed by heat fragmentation and cDNA synthesis. After enrichment, libraries were quantified using Quant-iT PicoGreen (Invitrogen, #P7589), normalized to 5 ng/µL, and pooled. The library pool was quantified using the KAPA Library Quantification Kit. All steps were performed in 96-well format with liquid handling by Agilent Bravo or Hamilton Starlet instruments. Pooled libraries were normalized to 2 nM and denatured with 0.1 N NaOH prior to sequencing. Flowcell cluster amplification and sequencing were performed on a NovaSeq 6000 (Illumina) according to manufacturer’s protocols, generating 2×151 bp paired-end reads with an eight-base index barcode read. DRAGEN was used for demultiplexing and data aggregation; all subsequent alignment and quantification steps were performed uniformly across both protocols on FASTQ files.

Reads were aligned to the appropriate reference genome using HISAT2 (v2.2.1) with splice-aware alignment enabled (--dta) and reverse-strand orientation specified (--rna-strandness R for single-end; RF for paired-end libraries). Resulting alignments were coordinate-sorted and indexed using SAMtools (v1.21). Gene-level read counts were quantified using featureCounts (Subread v2.0.1) by summarizing reads over exonic features aggregated by gene_id, with reverse-strand specificity (-s 2). All steps used Ensembl gene annotations matched to the appropriate genome assembly: Mus_musculus.GRCm38.102.gtf for mouse, Homo_sapiens.GRCh38.112.gtf for human, and Rattus_norvegicus.Rnor_6.0.102.gtf for rat.

For downstream analysis (TPM and differential expression), counts were restricted to protein-coding genes as defined by the respective Ensembl annotation. TPM values were calculated per sample for expression reporting, and differential expression analysis was performed using DESeq2 (v1.40.2) in R (v4.3.0), with genes requiring a minimum of 10 counts in at least 3 samples prior to analysis. Pairwise comparisons were evaluated using Wald tests, and genes were considered differentially expressed at an adjusted *p*-value < 0.01 (Benjamini–Hochberg) with log_2_ change thresholds specified in figure legends.

For cross-species gene comparisons, mouse and rat gene identifiers were mapped to their human orthologs using Ensembl BioMart via the R package biomaRt^74^. Mouse (*Mus musculus*, GRCm38) and rat (*Rattus norvegicus*, Rnor_6.0) datasets were aligned using Ensembl release 102 annotation, while human (*Homo sapiens*, GRCh38) datasets used Ensembl release 112. All human gene symbols were updated to current Ensembl 112 nomenclature by remapping through the human Ensembl gene ID after initial mapping. Genes lacking a one-to-one mapping were retained using their uppercase gene symbol as a fallback.

## Immunoblotting

Protein lysates for western blotting were prepared by washing cells in cold PBS and lysing in RIPA buffer (Boston BioProducts, #BP-115; 50 mM Tris-Cl, 150 mM NaCl, 0.50% sodium deoxycholate, 0.10% sodium dodecyl sulfate, 1% NP-40 substitute) supplemented with cOmplete protease inhibitor (Roche, #11836170001) and PhosSTOP phosphatase inhibitor (Roche, #4906845001). Samples were clarified by maximum-speed centrifugation at 4°C, and clarified lysate was normalized by BCA quantification.

The following antibodies were used: CREB1 (Cell Signaling, #9197), phospho-CREB1/ATF1 (Cell Signaling, #9198), CRTC3 (Cell Signaling #2720), CRTC2 (Bethyl Laboratories, #A300-637A), GAPDH (Cell Signaling, #2118), DYKDDDDK Tag (Cell Signaling, #14793), HA Epitope Tag (Cell Signaling, #3724), Anti-Rabbit HRP linked secondary (Cell Signaling #7074).

## CRTC2 CREB1-Binding Domain TurboID Labeling

1 × 10^6^ HEK293T cells were co-transfected with 1 µg of CRTC2^1–70–mTurbo–NLS–FLAG^combined with 1 µg of HA-ICER1 or 1 µg FLAG-mCherry-NLS. 24 hours after transfection, 50 µM biotin was added to the medium for 2 hours followed by a 1-hour washout in medium. Cells were lysed in RIPA buffer supplemented with protease and phosphatase inhibitors, and lysates were clarified by centrifugation before being bound to 50 µL magnetic streptavidin beads (Invitrogen, #65001) for 1 hour with rocking. Beads were washed four times in 1 mL RIPA, with the penultimate wash supplemented with an additional 500 mM NaCl. Protein was eluted from beads by incubation at 95°C for 5 min in β-mercaptoethanol-supplemented sample loading buffer and analyzed by western blotting. Co-transfected FLAG-tagged constructs were discriminated by size.

## Luciferase assay

4 x 10^5^ B16F10 cells were plated in a 24-well plate and incubated overnight. Cells were transfected the next day with 25 ng pCRE-Luc (4xCRE-HSV driven firefly luciferase construct, Stratagene), 25 ng pRL-CMV (Promega, E2261), and 400 ng empty plasmid DNA. The next day, cells were treated with DMSO or forskolin for 6 hr and lysed and processed with a Dual Luciferase kit (Promega, E1910), normalizing the pCRE firefly luciferin signal to the pRL-CMV coelenterazine signal for each sample before calculating the fold change of pCRE signal with forskolin over DMSO control.

## Guide RNA Sequences Human sgRNAs

sgCRTC3: gAGGGAAAACAGACGGAGCGA

sgICER1: GCTGTAACTGGAGATGACAC

sgAAVS1: GGGGCCACTAGGGACAGGAT

## Mouse sgRNAs

sgCrtc1: GCACAACCAGAAGCAGGCGG

sgCrtc2: GCAGAAGCAGCGTCAGGCCG

sgCrtc3: GCAGGGGTGCTGAGTAGTGG

sgCreb1: gCTATTAAATCACATACCTGT

sgAtf1: GTCATCGTTATCAGAAAGTG

sgCrem: GTTCCTACTCTAGCTCAGGT

sgCrem_all_isoforms: gAGAAGAAGCAACTCGCAAGC

## RT-qPCR Primer Sequences

### Human primers

MITF-M_F: CATTGTTATGCTGGAAATGCTAGAA

MITF-M_R: GGCTTGCTGTATGTGGTACTTGG

NR4A2_F: AAACTGCCCAGTGGACAAGCGT

NR4A2_R: GCTCTTCGGTTTCGAGGGCAAA

IL10_F: TCTCCGAGATGCCTTCAGCAGA

IL10_R: TCAGACAAGGCTTGGCAACCCA

SIK1_F: GCTCAAGGAGTATCGGAATGCC

SIK1_R: GTGGAAAGACCTTTCCTGAGGCA

OCA2_F: AGGAGAAGCGAGCACTCAGTGA

OCA2_R: CACCTGGGTTTCTACACTTCCG

TYRP1_F: TCTCAATGGCGAGTGGTCTGTG

TYRP1_R: CCTGTGGTTCAGGAAGACGTTG

DCT_F: CTCAGACCAACTTGGCTACAGC

DCT_R: CAACCAAAGCCACCAGTGTTCC

RPL11_F: GTTGGGGAGAGTGGAGACAG

RPL11_R: TGCCAAAGGATCTGACAGTG

### Mouse primers

Fos_F: GGGAATGGTGAAGACCGTGTCA

Fos_R: GCAGCCATCTTATTCCGTTCCC

Mitf-M_F: GCCTGAAACCTTGCTATGCTGGAA

Mitf-M_R: AAGGTACTGCTTTACCTGGTGCCT

Sik1_F: GCGACTACAACGAACAGGTGCT

Sik1_R: GTAGGAGGTAGTAAATGGCGGC

Oca2_F: CCGAGACAAGTCCTCATTGCAG

Oca2_R: GCTGTGAAACCAGCGAAGTCCA

Tyrp1_F: AGCCACAGGATGTCACTCAGTG

Tyrp1_R: GCAGGGTCATATTTTCCCGTGG

Dct_F: GCAAGATTGCCTGTCTCTCCAG

Dct_R: CTTGAGAGTCCAGTGTTCCGTC

Mef2c_F: GTGGTTTCCGTAGCAACTCCTAC

Mef2c_R: GGCAGTGTTGAAGCCAGACAGA

Trpm1_F: TGAGCAAGCGGAGGCTGATAAG

Trpm1_R: GCATCTTTGAGAGCCTCGGAGT

Gapdh_F: CTCCCACTCTTCCACCTTCG

Gapdh_R: GCCTCTCTTGCTCAGTGTCC

ICER_exon1_F: GCAAAAGCCCAACATGGCTGTAACTGGA

ICER_exon1_R: AAGTTGGCATGTCACCTGTGGCAGc

siRNA knockdown pools:

Human CRTC2: L-018947-00-0005

Mouse Crtc3: L-065875-01-0005

Mouse Crtc2: L-049303-00-0005

Non-targeting siRNA pool #2: D-001206-14-05

---

**Supplemental Figure 1.**
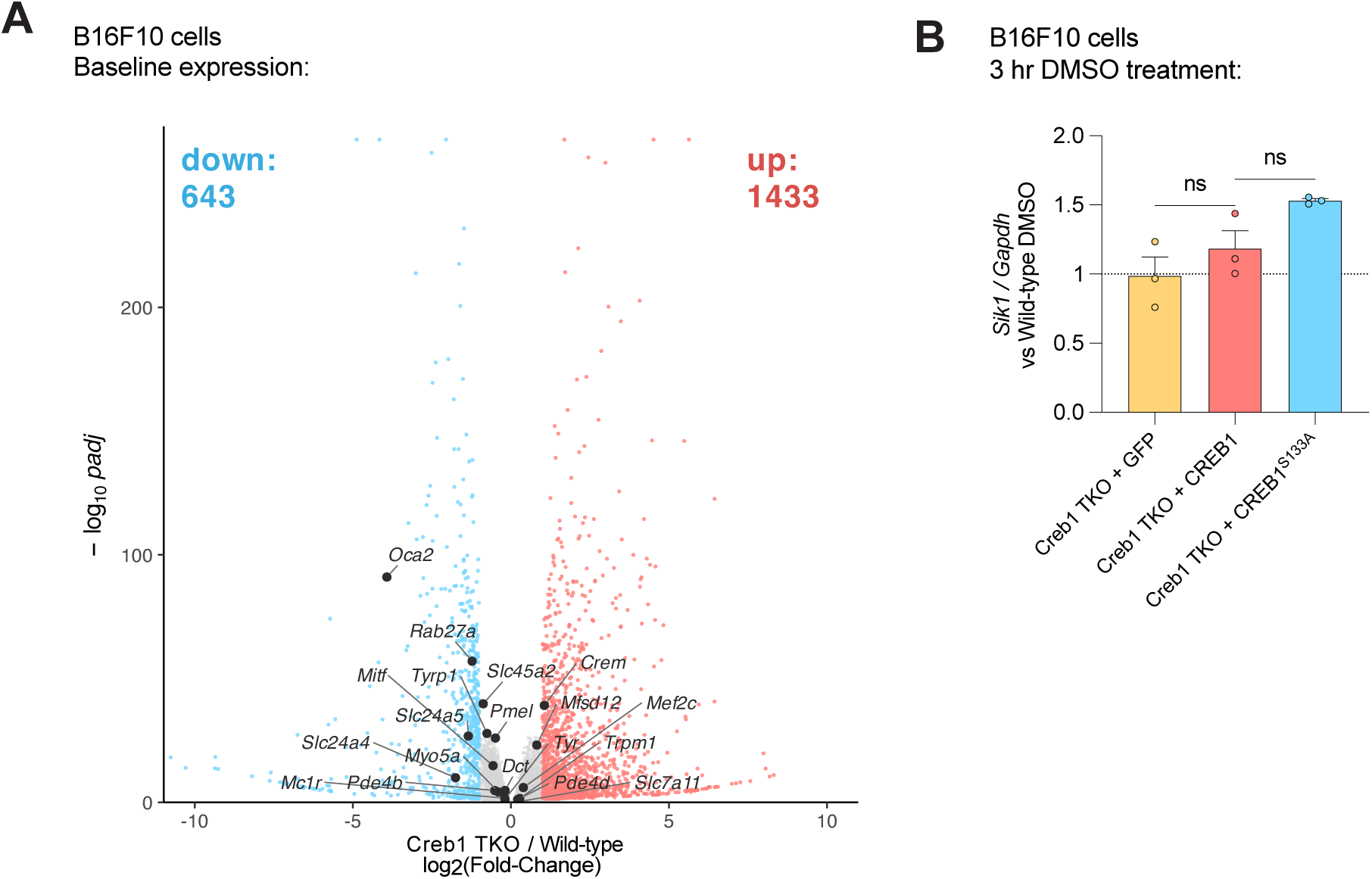
C**R**EB1 **paralog independent and dependent features of cAMP target gene regulation, related to Figure 1. (A)** Volcano plot comparing Creb1 TKO and wild-type cells from 3 hr DMSO treatment arms (“baseline”). Select pigmentation genes are highlighted. Analysis was restricted to protein-coding mRNAs, fold-change and adjusted p-values were calculated using DESeq2. **(B)** Baseline *Sik1* levels as measured by RT-qPCR and normalized to DMSO-treated wild-type cells. These data were collected alongside Figure 1E. *p<0.05, **p<0.01. Bar plot statistical tests were one-sample Student’s t-test against 1 (B). Error bars are standard error of the mean.

**Supplemental Figure 2.**
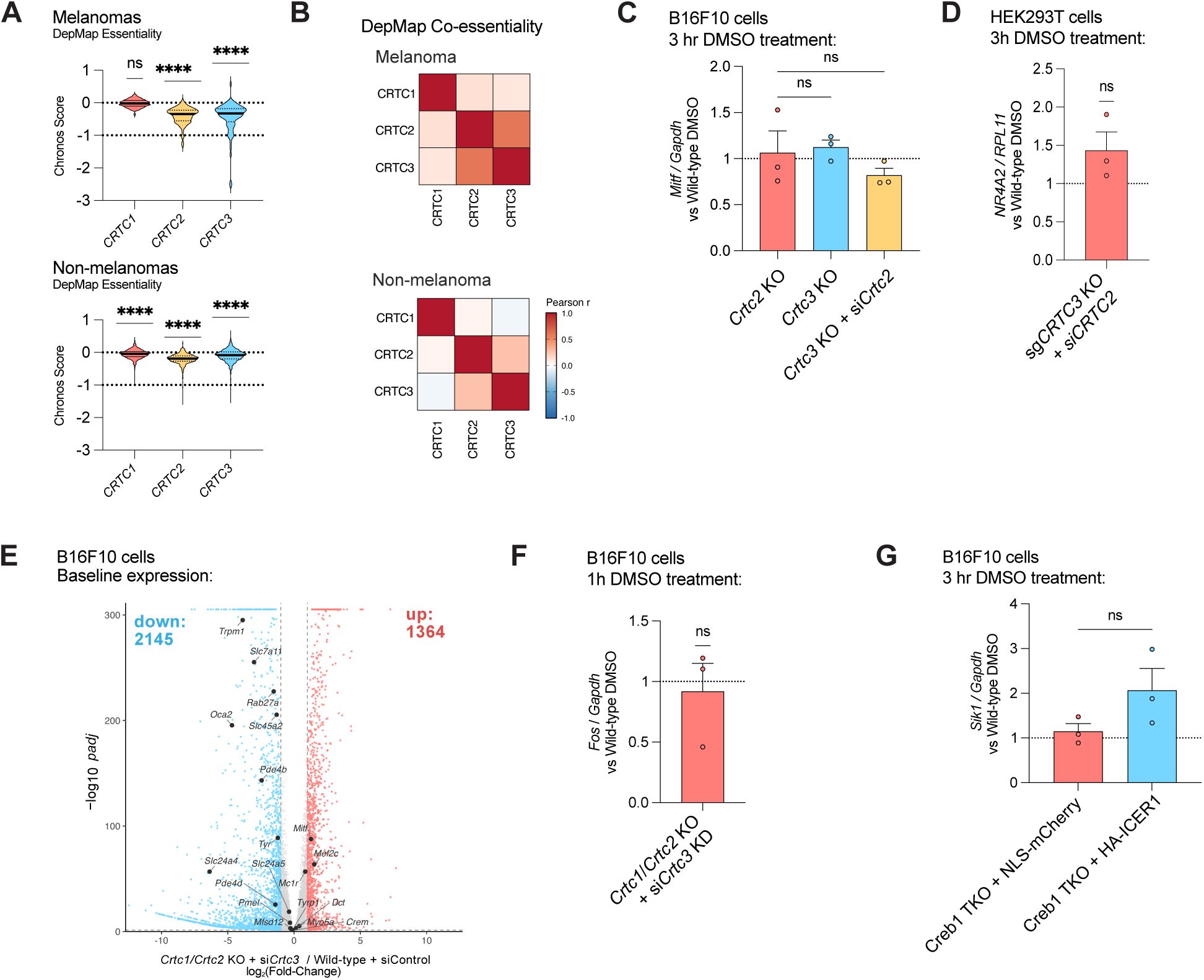
E**f**fects **of compound CRTC loss on baseline and cAMP-regulated gene expression, related to Figures 2 and 3. (A)** Chronos gene essentiality scores from DepMap 26Q1 for *CRTC* paralogs (melanoma, n = 80; non-melanoma, n = 1,178). **(B)** Correlation matrix (Pearson correlation) comparing Chronos gene essentiality scores between melanoma and non-melanoma cell lines (melanoma, n = 80; non-melanoma, n = 1,178). **(C)** Baseline *Mitf* levels as measured by RT-qPCR and normalized to DMSO-treated wild-type cells. These data were collected alongside Figure 2C (n = 3). **(D)** Baseline *NR4A2* levels as measured by RT-qPCR and normalized to DMSO-treated wild-type cells. These data were collected alongside Figure 2D (n = 3). **(E)** Volcano plot comparing Crtc1/Crtc2 knockout + *siCrtc3* to wild-type + siControl cells from 3 hr DMSO treatment arms (“baseline”). Select pigmentation genes are highlighted. Analysis was restricted to protein-coding mRNAs, fold-change and adjusted p-values were calculated using DESeq2. **(F)** Baseline *Fos* levels as measured by RT-qPCR and normalized to DMSO-treated wild-type cells. These data were collected alongside Figure 2F (n = 3). **(G)** Baseline *Sik1* levels as measured by RT-qPCR and normalized to DMSO-treated NLS-mCherry expressing cells. These data were collected alongside Figure 3C (n = 3). *p<0.05, **p<0.01, ***p<0.001, ****p<0.0001. Bar plot statistical tests were from ordinary one-way ANOVA with Sidak’s multiple comparison test (C, G) or one-sample Student’s t-test against 1 (D, F). Error bars are standard error of the mean.

**Supplemental Figure 3.**
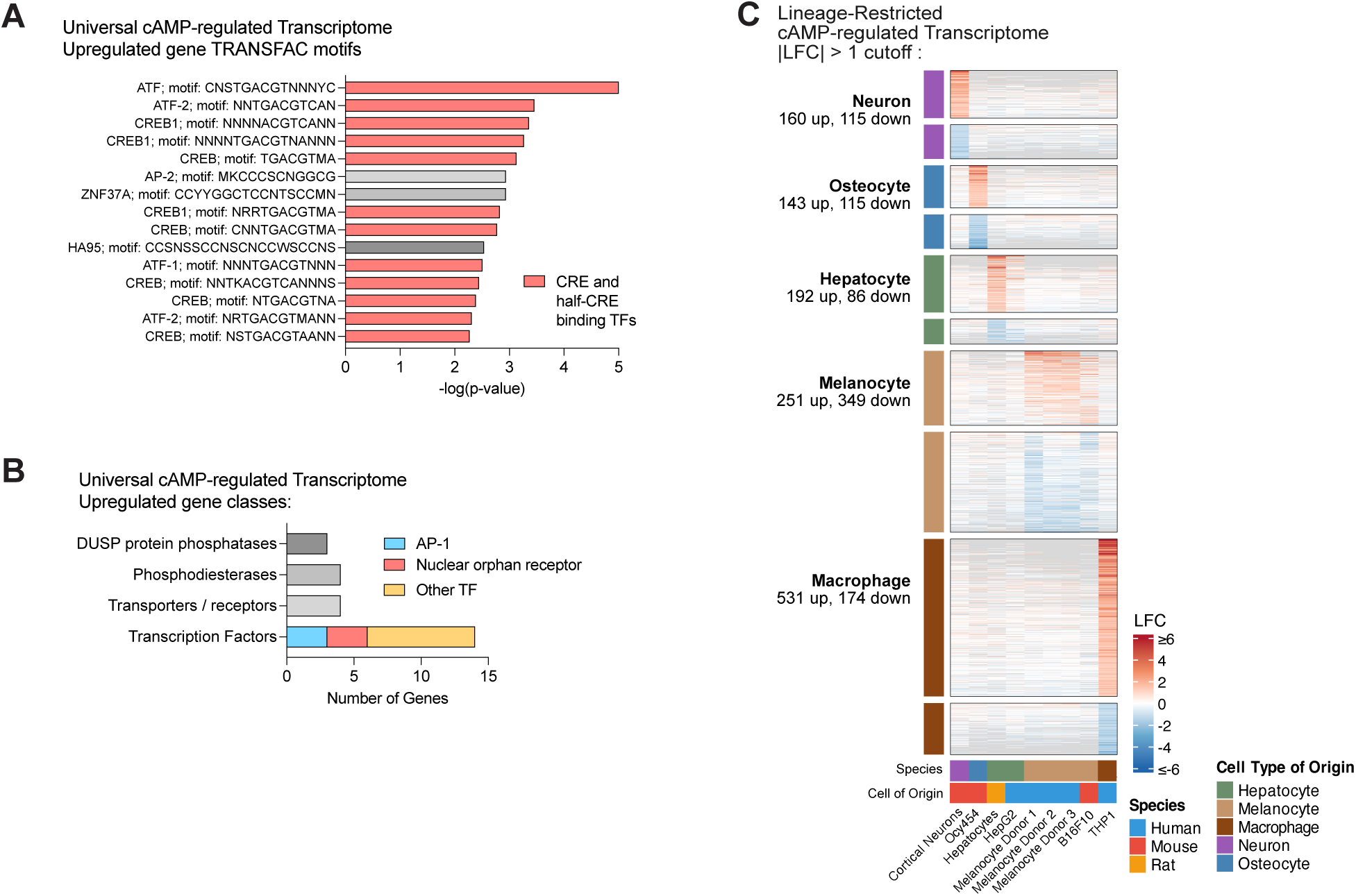
F**e**atures **of the universal and lineage-restricted cAMP-regulated transcriptional programs, related to Figure 4. (A)** TRANSFAC motif enrichment analysis from gProfiler analyzing upregulated genes from the UCAR (Figure 4A)^75^. The top 15 factors and their motifs are shown, 12/15 bind to CRE or half-CRE sequences (red columns). **(B)** Classification of upregulated genes from the UCAR (Figure 4A) highlighting the high number of transcription factors and other signaling regulators controlled by CREB1 in many cell types. **(C)** Lineage-restricted cAMP-regulated transcriptomes with a |LFC| > 1 and *padj* < 0.01 cutoff. An analogous plot with increased stringency (LFC| > 2, *padj* <0.01) is presented in Figure 4B.

**Supplemental Figure 4.**
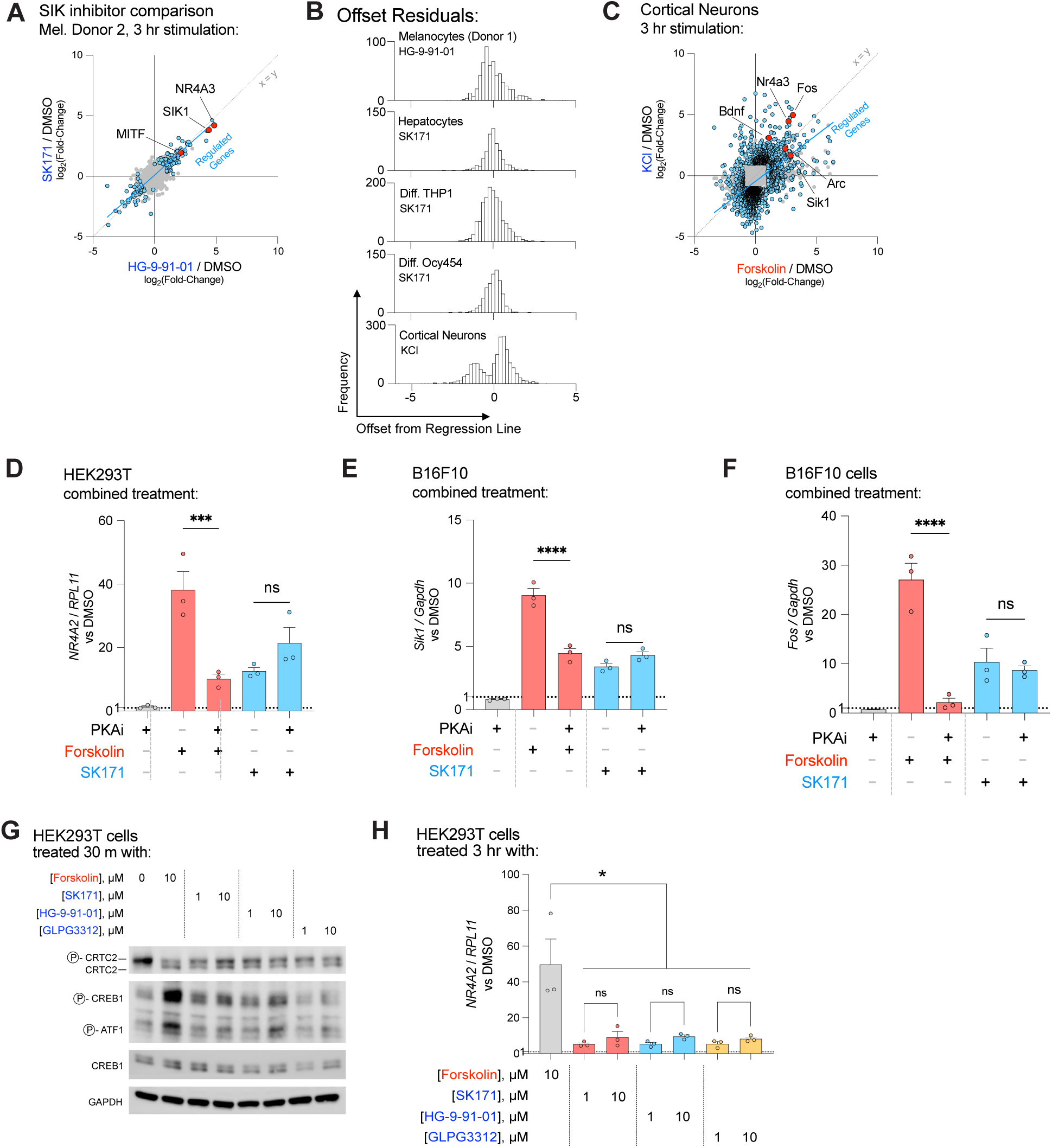
F**e**atures **of SIK inhibition compared to forskolin and cAMP-activation, related to Figure 5. (A)** Scatterplot of genes from melanocyte donor 2 cells treated with HG-9-91-01 or SK171 (1 µM each), demonstrating strong agreement between the two SIK inhibitors. Points and linear fit as in Figure 5B. **(B)** Histogram of perpendicular offsets of differentially regulated genes versus their regression line. Cell line and treatment (compared to forskolin) are labeled on inset. **(C)** Scatterplot of genes from neuronal cells treated with forskolin or KCl based depolarization. Points and linear fit as in (A). **(D-E)** *NR4A2* and *Sik1* induction measured by RT-qPCR from HEK293T and B16F10 cells, respectively. Cells were pretreated for 1 hr with the PKA inhibitor BLU0588 (2 µM) before a 3 hr treatment with forskolin (10 µM) or SK171 (1 µM). Samples were normalized to *RPL11* or *Gapdh* and DMSO-treated cells (n =3). **(F)** *Fos* induction measured by RT-qPCR from B16F10 cells. Cells were pretreated for 1 hr with the PKA inhibitor BLU0588 (2 µM) before a 1 hr treatment with forskolin (10 µM) or SK171 (1 µM). Samples were normalized to *Gapdh* and DMSO-treated cells (n =3). **(G)** Immunoblot of HEK293T cells treated for 30 min with forskolin (10 µM) or the SIK inhibitors SK171, HG-9-91-01, or GLPG3312 (1 µM and 10 µM). **(H)** *NR4A2* induction measured by RT-qPCR from HEK293T cells, normalized to *RPL11* and DMSO control (n = 3). *p<0.05, **p<0.01, ***p<0.001, ****p<0.0001. Bar plot statistical tests were from ordinary one-way ANOVA with Sidak’s multiple comparison test. Error bars are standard error of the mean.

